# Age-related cerebello-thalamo-cortical white matter degradation and executive function performance across the lifespan

**DOI:** 10.1101/2025.08.22.671779

**Authors:** Jessica N. Kraft, Alyssa Ortega, David A. Hoagey, Karen M. Rodrigue, Kristen M. Kennedy

## Abstract

The cerebellum supports higher-order cognition, such as working memory and executive function (EF) both directly and through connection with prefrontal areas via cortical loops. Thus, age-related degradation to white matter connectivity comprising cerebello-thalamo-cortical (CTC) loops may underlie age-related differences in EF. In 190 healthy adults (aged 20-94 years) we collected diffusion tensor imaging scans and multiple tests of working memory and EF. Deterministic tractography was used to generate CTC tracts from which white matter metrics (mean, radial, axial diffusivities) were extracted. General linear model results indicated that reduced white matter integrity (i.e., higher diffusivity) was associated with significantly poorer EF performance in an age-dependent fashion. Higher mean, radial, and axial diffusivities in fronto-cerebellar white matter was associated with lower EF scores in older, but not younger, adults. These findings suggest CTC white matter connectivity is important for executive function performance and lend mechanistic evidence to the role of the cerebellum in age-related differences in higher-order cognitive operations.

## INTRODUCTION

The cerebellum plays a critical role in cognition, along with its well-established role in motor coordination. Three decades of research has implicated the cerebellum in cognition and affect, beginning with clinical observations of cerebellar lesions with no motor phenotypes, but marked deficits in executive function, language processing, visuospatial performance and dysregulation of affect, termed cognitive affective syndrome (CCAS) (Argyropoulos et al., 2020; J. D. Schmahmann, 1991; J. D. Schmahmann & Sherman, 1997). Such findings provided the impetus to explore cerebellar contributions to non-motor function. Much as lesions to the anterior lobe of the cerebellum cause dysmetric motion, lesions to the posterior cerebellum result in deficits of “dysmetria of thought.” (J. D. Schmahmann, 1991, 1996, 2019)

### The Structure of the Cerebellum

The cerebellum is a remarkably complex structure. Recent estimates suggest that it contains between 50-80% of the brain’s total number of neurons, and is about 78% of the size of the cerebrum in surface area, despite comprising only 10% of the brain’s total volume (Herculano-Houzel, 2009, 2012; Lyu et al., 2024; Silver, 2018). The cerebellum is parsed into ten lobules contained within three lobes; lobules I through V comprise the anterior lobe, lobules VI through IX make up the posterior lobe, and lobule X is known as the flocculonodular lobe (J. D. Schmahmann, 2019; Stoodley et al., 2012). Lobule VII, the largest in the human brain, is further segregated into three sections-Crus I, Crus II and area VIIB (Lyu et al., 2024; J. D. Schmahmann, 2019). Within the white matter of the cerebellum lie the deep cerebellar nuclei (the fastigial, interposed, and dentate nuclei), through which all efferent connections travel to cortical regions (Diedrichsen et al., 2011; Küper et al., 2012) The anterior cerebellum (particularly lobules IV to VI), combined with portions of lobule VIII, has dense projections to the primary motor cortex and premotor areas (Middleton & Strick, 2000; Palesi et al., 2015; Stoodley & Schmahmann, 2009; Strick et al., 2009). Conversely, the posterior cerebellum, namely Crus I, Crus II and area VIIB has strong connections to the prefrontal cortex, with few connections to cortical sensorimotor regions (Argyropoulos et al., 2020; Lyu et al., 2024; J. D. Schmahmann, 2019; Stoodley et al., 2012; Strick et al., 2009).

The distinct cortical connections of the anterior and posterior cerebellum lend itself to the formation of discrete corticocerebellar loops. As its name implies, the afferent cortico-ponto-cerebellar (CPC) pathway commences at the cerebral cortex, synapses in the anterior pontine nuclei, and decussate prior to the contralateral middle cerebellar peduncle before reaching the cerebellar lobules. In contrast, the efferent cerebello-thalamo-cortical (CTC) pathway, originating at the cerebellum, travels through the ipsilateral dentate nucleus, ascends the superior cerebellar peduncles and through the contralateral anterior and lateral thalamic nuclei to the cerebral cortex (Palesi et al., 2015, 2017; Ramnani et al., 2006)(for a review, see (D’Angelo, 2018). The CTC ascends as far as the frontal lobe, with more fibers terminating in the prefrontal cortex than any other region of the cerebrum (Palesi et al., 2015). The closed loop circuitry of the CTC pathway provides insight into the well-established role of the cerebellum in cognitive performance.

### The Cerebellum and Cognition

The convergence of structural and functional neuroimaging studies has long demonstrated that the cerebellum is essential in multiple domains of cognition. Posterior cerebellar activation is widely observed during tasks of working memory, executive functioning, and language production, outside of motor response preparation (Desmond et al., 1997; Guell & Schmahmann, 2020; J. D. Schmahmann, 2019; Tomasi et al., 2007). Advancements in functional neuroimaging have demonstrated that various cognitive domains are represented throughout the posterior cerebellum with topographic specificity. A meta-analysis by Stoodley and Schmahmann (2009) found that peak activation during tasks of language and verbal working memory were within the junction of lobule VI and Crus I, whereas executive function tasks activated the majority of Crus I, as well as left lobules VI and VIIB. Furthermore, this topography representation occurs even within a single cognitive domain, based on functional task features. For instance, the strength and location of cerebellar activation during working memory tasks was dependent on the stimuli used, with numerical working memory stimuli lateralized to left Crus I and verbal working memory stimuli lateralized to right Crus I (King et al., 2019).

Perhaps unsurprisingly, structural and functional declines to the cerebellum and associated areas (e.g. through aging process) are implicated in decreased cognitive performance. Working memory has shown to be significantly associated with left posterior cerebellar volume, while reaction time was related to Crus I specifically (Bernard & Seidler, 2013). Studies examining cerebellar activation provide empirical support that the cerebellum is integrally linked with cognitive control regions, and that there are functionally relevant age-related differences in this connectivity. Specifically, the strength of resting-state cortico-cerebellar connections is weaker and more widespread in older adults, compared to younger adults; moreover, this differential pattern of functional connectivity was associated with global levels of cognition, above and beyond the influence of age alone (Bernard et al., 2013, 2020; Bernard & Seidler, 2013; Hausman et al., 2019). Task-based functional MRI studies have shown co-activation of lobule VII and prefrontal cortices during completion of both visual and verbal working memory tasks (Chen & Desmond, 2005; Hayter et al., 2007).

Deficits in executive functioning and visual spatial processing are hallmark characteristics of CCAS, suggesting that the cerebellum is heavily involved in these intricate cognitive processes (Argyropoulos et al., 2020; J. D. Schmahmann, 2019; J. D. Schmahmann & Sherman, 1997; Stoodley & Schmahmann, 2009). Task-based functional connectivity analyses have demonstrated error-related activation in the thalamus and cerebellum during completion of executive functioning tasks (Ide & Li, 2011). An activation likelihood estimation (ALE) meta-analysis of cerebellar activation of over 700 healthy participants found non-overlapping activation peaks within the cerebellum associated with both executive functioning and working memory. Specifically, regions within the left lobule VI, left Crus I, right Crus I and left VIIB were activated during executive functioning, whereas regions associated with working memory were mainly right lateralized and included the right lobule VI, right VIIIA, right lobule VI, and right Crus I (Stoodley & Schmahmann, 2009). In regions of overlapping cerebellar activation, peaks were significantly stronger in the left lobule VI, left Crus I, and left VIIB during completion of executive functioning tasks, compared to working memory tasks (Stoodley & Schmahmann, 2009). Furthermore, a recent study using q-space diffeomorphic reconstruction found that quantitative anisotropy (QA) values between the cerebellum and fornix were associated with performance on tasks of working memory and inhibitory control (Parsaei et al., 2025).

### Brain Aging and the Cerebellum

Brain aging is regionally heterochronic; certain regions of the brain, such as the dorsolateral prefrontal cortex (DLPFC), basal ganglia, hippocampus and cerebellum are more vulnerable to the effects of aging (Jernigan et al., 2001; Raz et al., 2004, 2010). Volumetric declines of cerebellar grey matter throughout the aging process are significant, and indeed are comparable to declines in prefrontal cortices (G. E. Alexander et al., 2006; Bernard & Seidler, 2013; Hoogendam et al., 2012; Jernigan et al., 2001; Raz et al., 2010). Of potential greater concern is that white matter volumetric reductions in the cerebellum are even steeper than those of grey matter, with the highest declines occurring in Crus I and Crus II of the cerebellum (Abe et al., 2008; Hoogendam et al., 2012; Raz et al., 2010, 2013).

Concomitant with volumetric white matter declines in the cerebellum, research has suggested that the integrity of afferent and efferent white matter tracts in the cerebellum may be susceptible to age-related decline as well. Diffusion tensor imaging (DTI), an indirect method of assessing white matter structural integrity, quantified most frequently by four metrics. The two most common are fractional anisotropy (FA) and mean diffusivity, which measures the directionality of diffusion and magnitude of diffusion, respectively. Additional metrics can be used to further quantify the shape of a diffusion tensor, such as axial diffusivity (AD) which is a measure of the largest eigenvector in a tensor, and radial diffusivity (RD) which is an average of the remaining two shortest tensors (A. L. Alexander et al., 2007; O’Donnell & Westin, 2011). Several DTI studies have found that age was associated with lower FA values and higher RD/AD/MD values in cerebellar white matter and cerebellar peduncles, suggesting poorer white matter integrity inside the cerebellum with increasing age (Bennett et al., 2010; Cavallari et al., 2013; Kafri et al., 2013).

Considering the neurophysiology of the CTC pathway, findings of age-related degradation of white matter in the cerebellum and adjoining structures, as well as reported associations between cerebellar structure and function with working memory and executive function performance, the aims of the current study were to generate reliable white matter tracts forming the CTC pathway across the adult lifespan, and assess the vulnerability of the integrity of this white matter tract to lifespan aging. Furthermore, the current study aims to assess whether the CTC was associated with cognitive performance on a battery of executive functioning and working memory tasks, and if this association was moderated by age. We hypothesized that increasing age would be associated with higher cerebello-thalamo-fronto white matter diffusivity metrics (MD, RD and AD), and that poorer white matter integrity in the CTC pathway would be associated with poorer performance on working memory and executive function tasks. Further, we hypothesized that these white matter-cognition associations may be age-dependent, occurring only by middle or older age.

## METHODS

### Participants

As part of our ongoing longitudinal study (Dallas Area Longitudinal Lifespan Study of Aging; D.A.L.L.A.S.), 190 cognitively normal adults with complete cognitive and MRI data were selected for the current study (mean age = 53.66 ± 18.87, age range 20-94). Participants had no history of cardiovascular, neurologic, psychiatric, or metabolic problems, head trauma with loss of consciousness, or substance abuse. All participants were fluent English speakers, right-handed, passed hearing acuity screening, and had corrected-to-normal vision. Participants were excluded at the in-person screening visit if they met criteria for possible depression or dementia, using cutoff criteria from the Center for Epidemiologic Studies Depression Scale (CES-D) scores > 16 (Radloff, 1977) and Mini Mental State Exam (MMSE) scores ≤ 26 (Folstein et al., 1975), respectively. MRI scanning took place on average within 50 days of neuropsychological data acquisition (M = 50.04, SD = 41.72 days). Table 1 reports demographic information, as well as cognitive composite scores. All participants provided written informed consent in accordance with the local Institutional Review Boards.

**Table 1.** Sample Demographics.

| **Demographic characteristics** |  |
| --- | --- |
| Number of participants | 190 |
| Age, mean ± SD | $53.66 \pm 18.87$ |
| Sex, N (%) |  |
| Men | 78 (41.05%) |
| Women | 112 (58.95%) |
| Years of Education, mean ± SD | 15.52 ± 2.47 |
| Category Switching Task Total, mean ± SD | 14.51 ± 3.17 |
| Category Switching Task Correct, mean ± SD | 13.40 ± 3.35 |
| Trail Making Task (sec), mean ± SD | 79.14 ± 36.46 |
| Color Word Interference (sec), mean ± SD | 60.25 ± 17.17 |
| LSPAN Simple Score, mean ± SD | 3.19 ± 1.11 |
| LSPAN Absolute Score, mean ± SD | 21.90 ± 12.35 |
| LSPAN Total Score, mean ± SD | 51.87 ± 13.31 |
| WAIS Digit Span Forward, mean ± SD | 11.12 ± 2.30 |
| WAIS Digit Span Backwards, mean ± SD | 9.22 ± 2.54 |
| WAIS Digit Span Sequencing, mean ± SD | 9.18 ± 2.10 |

### Neuroimaging Acquisition Protocol

All neuroimaging data were acquired with a 3T Philips Achieva scanner with 32-channel head coil using SENSE encoding at the University of Texas Southwestern Advanced Imaging Research Center. Diffusion weighted images were collected using a single shot echo-planar sequence with 65 whole brain axial slices, 30 diffusion-weighted gradient directions, b = 1000, 1 non-diffusion weighted b0, TR 5611 ms, TE = 51 ms, flip angle = 90°, voxel size = 2mm isotropic, slice thickness = 2.2 mm, acquisition time = 4:19 minutes. T1-MPRAGE images were collected with 160 sagittal slices, voxel size 1mm isotropic, flip angle=12°, TR/TE = 8.1ms/3.7ms, FOV=256 x 204 x 160, matrix=256×256, acquisition time = 3:57 minutes.

### MRI Data Processing and Tractography Protocol

T1 images were skull stripped via BET (Smith, 2002), intensity bias corrected, and registered to Montreal Neurological Institute 1 mm template space (Montreal Neurological Institute, McGill University, Canada) using ANTS (Avants et al., 2011). All diffusion images were processed and quality control checked using DTIPrep v1.2.4 to identify potential susceptibility distortions or eddy current artifacts (Oguz et al., 2014). Raw DWI volumes were denoised and gradients containing substantial intensity or movement distortions were detected and removed from subsequent analyses. Remaining gradients were corrected using the default parameters in DTIPrep and co-registered to the non-diffusion weighted b0 image. Following artifact correction, diffusion directions were adjusted to account for reorientation of individual gradients and to preserve the original encoding direction (Leemans & Jones, 2009). Deterministic tractography of cerebello-thalamo-cortical (CTC) tracts and calculation of white matter diffusivities (AD, RD and MD) was performed using DSI Studio 3 (Yeh et al., 2013).

### Regions of Interest (ROI) and Regions of Avoidance (ROA)

Frontal ROIs were comprised of bilateral 12 mm spherical ROIs covering the anterior limb of the internal capsule, near the genu of the corpus callosum. The left frontal ROI was centered at MNI x = -18, y = 15, z = 2 and the right frontal ROI was centered at x = 17, y = 15, z = 2. Frontal ROIs were then registered to native space using a series of non-linear transformations via ANTS for each participant. The cerebellar ROI was derived from individual subjects’ bilateral cerebellum parcellations via FreeSurfer v5.3.0 using the SUIT atlas (Diedrichsen, 2006). See Figure 1.

**Figure 1:**
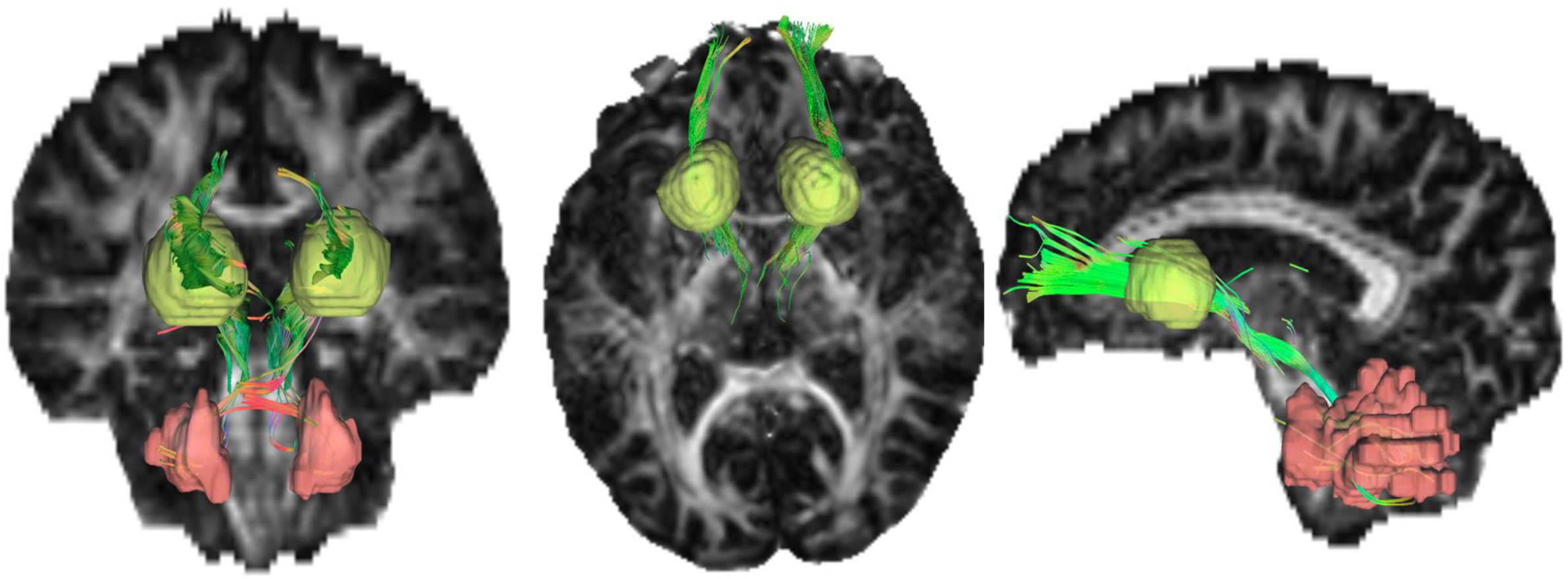
Cerebello-thalamo-cortical tract obtained from deterministic tractography from a representative participant overlaid on their native space FA map viewed from coronal, axial, and sagittal planes. Pink ROI is the 3D cerebellar hemispheres, yellow ROIs are the anterior limb of the internal capsule.

In addition, we defined 5 anatomical regions of avoidance (ROAs) to best isolate streamlines running through the prefrontal cortex, thalamus, and dentate nucleus of the cerebellum. One ROA consisted of an axial exclusionary plane at z = -56 to exclude spurious fibers projecting to the brainstem. A second ROA combined the premotor, primary motor, somatosensory cortex, and temporal pole (Brodmann areas 1-8 and 37-41) to restrict fibers to the prefrontal ROIs. The third ROA included the entire corpus callosum, to preserve hemispheric connections and to avoid inclusion of callosal fibers. The fourth ROA comprised a 10mm spherical ROA located at the pons, centered at MNI x=1, y=-22, z=-36 to eliminate fibers crossing between right and left hemispheres of the cerebellum. The final ROA consisted of bilateral 12 mm ROIs between the splenium of the corpus callosum and Brodmann area 41 (left ROA centered at MNI x=-26, y=-32, z=14 and right ROA centered at MNI x=24, y=-32, z=14) to eliminate spurious fibers that traveled to the inferior longitudinal fasciculus. All ROAs were created in MNI space and warped to native space via ANTS for each participant, followed by manual inspection of the ROIs. The following parameters were used in the deterministic tracking algorithm: maximum turning angle = 60 degrees, step size = 1 mm, minimum/maximum length = 20/500 mm, and FA threshold > 0.12. Diffusivities were extracted from tract voxels and multiplied by 1,000. Figure 1 illustrates the CTC streams obtained from deterministic tractography from a representative participant. Mean diffusion metric values (AD, MD, RD) were calculated within hemisphere and then averaged across both hemispheres to obtain one mean metric per participant.

### Cognitive Measures

#### Executive Function Composite

The executive function (EF) construct included scores from the Trail Making Test (Letter-Number Switching score), Verbal Fluency Task (Category Switching score), and Color-Word Interference Test (Inhibition/ Switching score) from the D-KEFS (Delis-Kaplan Executive Function System) cognitive battery (Delis et al., 2001). A standardized composite score was created to represent each participant’s EF ability by scaling measures so that higher scores reflect better performance across all tasks, standardizing measures to Z-scores for each task (mean of 0, SD 1), and averaging all measures for each participant.

#### Working Memory Composite

The working memory construct was comprised of the subscores of the Listening Span task (LSPAN) (Daneman & Carpenter, 1980) including simple span, absolute span and total span, as well as the sequencing, forward and backwards subtasks from the WAIS Digit Span test (WAIS IV)(Wechsler, 2008). As with the EF composite, all scores were scaled so that higher scores reflected better performance and then standardized to Z-scores and averaged across all measures for each participant.

### Statistical Analysis Approach

Analyses were conducted using the “lm” linear models function in the R statistical software package (R Core Team, 2014). Separate moderation models were tested to determine whether executive function or working memory performance (as dependent variables in separate models) varied differentially as a function of tract diffusivity metric (AD, RD, MD in separate models) and age (a continuous mean-centered predictor) or their interactions. Models also included sex and years of education as covariates. To test for nonlinear effects of age, age^2^ was entered into the models (along with linear age) and removed when non-significant. Significance level was set to *p* < .05 for all models. Effect sizes are expressed as semi-partial correlations (*sr^2^*), which express the unique contribution of each regression term on the dependent variable after controlling for all other terms in the model. Semi-partial correlation coefficients can be interpreted as r^2^ values for an individual predictor, and as such, semi-partial correlation coefficient of 0.01 is generally considered a small effect, 0.09 is considered medium and 0.25 or higher is considered a large effect (Cohen, 1988). Significant interaction terms containing continuous variables were decomposed using the Johnson-Neyman procedure with simple slopes estimation (Johnson & Neyman, 1936).

## RESULTS

### Effects of Age on CTC White Matter Diffusivity Metrics

Three general linear models were assessed, with diffusion metric as the dependent variable (MD, RD, or AD), age, age^2^ as independent variables, and sex and years of education as covariates. Significant main effects of quadratic age were found on all three diffusivity metrics: MD [F(1,185) = 23.232, p< 0.001, *sr^2^*=0.09], RD [F(1,185) = 8.898, p = 0.003, *sr^2^*= 0.04] and AD [F(1,185) = 51.782, p < 0.001, *sr^2^*= 0.17], describing an accelerated increase in diffusivity with increasing age (see Appendix Table 1 for regression model details). Figure 2 suggests this acceleration inflection point to be around approximately age 50.

**Figure 2:**
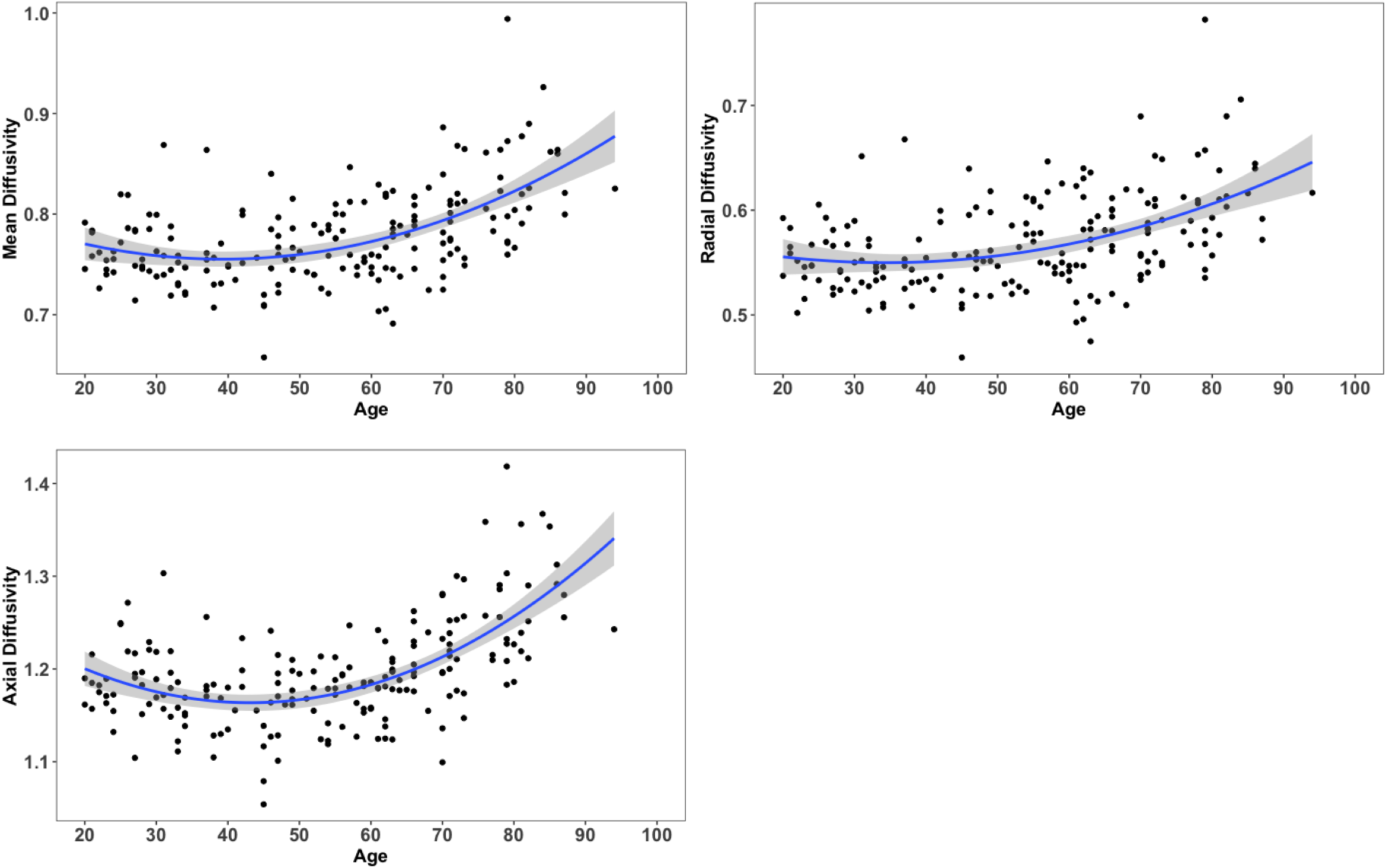
Effects of age on cerebello-thalamo-cortical tract mean, radial, and axial diffusivities. An accelerated rate of increased diffusivity with increasing age across the adult lifespan was found for all three white matter metrics.

### Effects of CTC White Matter Metrics on Executive Function (EF) Performance

We next aimed to assess whether age-related cerebello-frontal white matter properties were associated with executive function performance. Three linear models were run with EF composite score as the dependent variable and age, diffusivity (MD, RD, or AD) and their interactions as independent variables, with sex and years of education as covariates. We found significant age × diffusivity interactions on EF performance for all three metrics, indicating that the effects of tract diffusivity on EF were age-dependent: age × MD [F(1,184) = 6.027, p = 0.015, *sr^2^*= 0.02], age × RD [F(1,184) = 4.601, p = 0.033, *sr^2^*= 0.02], and age × AD [F(1,184) = 7.756, p = 0.006, *sr^2^*=0.03]. See Appendix Table 2 for additional model parameters.

To decompose these significant interactions, we used the Johnson-Neyman method with simple slopes for visualization. Slopes were estimated at three age levels for all significant interactions: -1 standard deviation (SD) below the mean (∼35 years old), at the mean age (∼54 years old) and + 1 SD above the mean (∼73 years old). Simple slopes analyses, illustrated in Figure 3, revealed that the moderating association between white matter diffusivity and executive function is primarily driven by older adults (blue lines) for each of the white matter metrics (i.e., those +1 SD above the mean). Johnson-Neyman interval calculations (right-side panels of Figure 3) estimate that the association between MD and EF performance becomes significant around age 58, between RD and EF performance around age 57, and between AD and EF performance around the age of 61 (panels a, b, and c, respectively).

**Figure 3:**
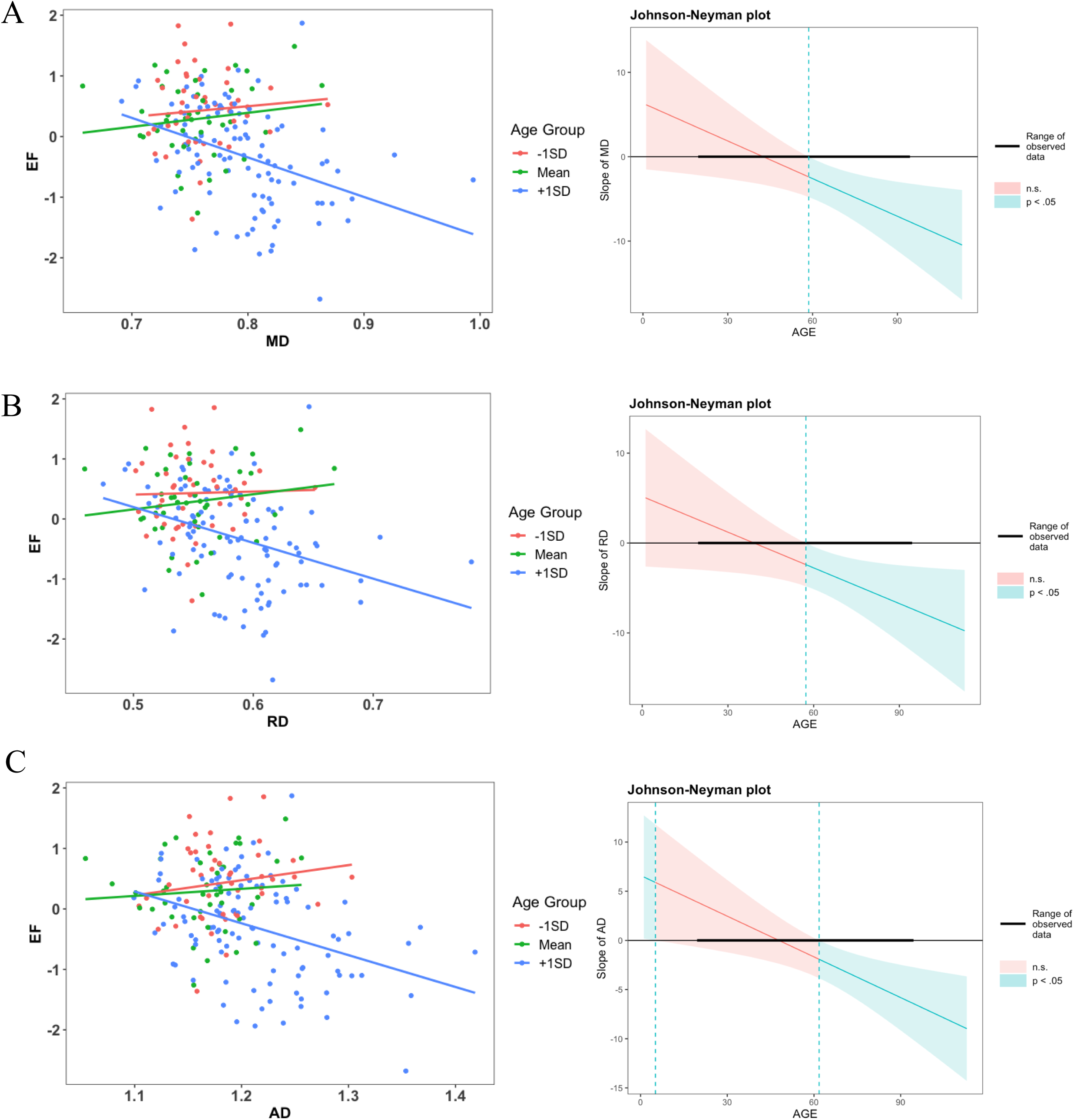
Age moderates the association between executive function scores and white matter integrity. Significant age × diffusivity interactions for MD (A), RD (B), and AD (C) on executive function performance are decomposed using simple slopes (left panels) and Johnson-Neyman interval ranges (right panels). The age-dependent diffusivity is due to the older adults, where this association becomes significant around late middle age (57-61 years depending on white matter metric).

#### Effects of CTC White Matter Metrics on Working Memory Performance

In contrast to executive function performance, there were no significant main effects of age, white matter metric (MD, RD, AD), and no significant age by white matter metric interactions on the working memory composite performance (all p-values > 0.378).

## DISCUSSION

The current study assessed cross-sectional age-related differences across the adult lifespan in cerebello-thalamo-cortical pathway white matter integrity as indexed by diffusivity metrics, and the association with cognition. We found an accelerated rate of decline of mean, radial, and axial diffusivity in the CTC tract, evidenced by significant main effects of quadratic age on all three diffusivity metrics. It is well established that white matter metrics decline with increasing age (Bennett & Madden, 2014; Kennedy & Raz, 2009; Madden et al., 2012; Raz & Rodrigue, 2006) and several studies have found an identical pattern of accelerated aging in MD, RD, and AD in other white matter regions of the brain (Damoiseaux et al., 2009; Sexton et al., 2014). The current study aligns with empirical support that the cerebellum exhibits some of the steepest age-related declines, comparable to those of the prefrontal cortices (Abe et al., 2008; G. E. Alexander et al., 2006; Bernard & Seidler, 2013; Raz et al., 2010). The present study expands these findings, demonstrating that white matter tracts bridging the cerebellum to the prefrontal cortex are also impacted in an age-dependent fashion.

Extant literature has further suggested that volumetric changes in the cerebellum are associated with cognitive performance throughout the aging process, which was partially supported in the current study. We found significant age by diffusivity metric interactions on EF, suggesting that increasing RD, MD, and AD is associated with worse EF performance. These findings are corroborated by previous research, demonstrating that BOLD activation throughout the CTC tract, including Crus I, Crus II and prefrontal cortices, is specifically associated with functional connectivity in the executive control network (Habas et al., 2009; Seeley et al., 2007). Functional activation of the cerebellum has been evidenced during completion of executive function tasks, including the Wisconsin Card Sorting Task (Lie et al., 2006), Tower of London (Schall et al., 2003), as well as the Stroop Inhibition task used in the current study (Ravnkilde et al., 2002). Recent work has further suggested that functional coherence between nodes throughout the CTC tract can reliably distinguish among healthy controls, individuals with mild cognitive impairment, and individuals with Alzheimer’s disease (Yao et al., 2025). Executive functions encompass several aspects of cognition, such as working memory, set shifting, inhibition and fluency (Rabinovici et al., 2015; Ribeiro et al., 2024). Nevertheless, research has delineated distinct regions of the cerebellum associated with canonical aspects of executive function (such as set shifting and response inhibition), while controlling for working memory, language and spatial processing (Stoodley & Schmahmann, 2009). Given the cerebellum’s susceptibility to structural degradation in aging (Kennedy & Raz, 2009), its pivotal role in executive functioning (Baumann et al., 2015; Koziol et al., 2014) and its extensive connections to prefrontal cortices also associated with executive functioning (Palesi et al., 2015), integrity of the CTC tract appears to be beneficial for executive functioning in older age.

Contrary to what might be expected from the literature, we did not find a significant relationship between diffusion metrics of the CTC pathway and working memory performance. This is surprising, as a substantial body of work has demonstrated the relationship between working memory and functional activation of cerebellar lobules (Bernard & Seidler, 2013; Chen & Desmond, 2005; Koziol et al., 2014; Stoodley et al., 2012; Stoodley & Schmahmann, 2009). It may be that white matter degradation in the CTC tract differentially impacts performance on set-shifting and inhibition tasks, compared to working memory paradigms. Studies have shown that subcortical structures, including the thalamus, are essential in executive control (Baumann et al., 2015; Rabinovici et al., 2015; J. Schmahmann & Pandya, 2008). A large body of literature has demonstrated that executive functioning performance and white matter integrity are heavily correlated and reliably established in older age (Bennett & Madden, 2014; Gustavson et al., 2023; Madden et al., 2004; Ribeiro et al., 2024) whereas the relationship between working memory and white matter integrity, while important, is less clear (for review see Lazar, 2017). An alternative possibility may be due to the duration of delay with the tasks. Much of the literature on working memory and the cerebellum employs tasks with longer delays than the working memory tasks in our composite. For instance, the Sternberg Working Memory paradigm requires participants to encode a series of stimuli and maintain that information until the retrieval phase (Sternberg, 1966). Indeed, studies have found that performance on this task is associated with the cerebellum (Cairo et al., 2004; Chen & Desmond, 2005; Kirschen et al., 2005). However, upon shortening the working memory delay in the Sternberg task (Chen & Desmond, 2005) and a similar verbal working memory task (Bohland & Guenther, 2006), activation in the cerebellum was no longer observed, compared to longer delays of the same task. It is possible that activation of the cerebellum in working memory is dependent on the duration of working memory delays, however, more work should be done to investigate this association.

While the current study provides valuable insights into the role of the CTC tract in cognitive performance, it is not without its limitations. First, the study utilized a cross-sectional design, and therefore cannot estimate within-person change effects. Future longitudinal studies measuring change in CTC tract properties and cognitive performance over time are needed. Second, as suggested above, the working memory paradigms used in the current study utilized short delays, which were not altered based on accuracy. Future research should investigate the role of the cerebellum in the context of varying working memory delays and difficulty levels and may find more age- or white matter sensitive outcomes. Third, executive function as a construct involves multiple lower-level cognitive processes, such as attention, set-shifting, reasoning and decision making, which are inherently associated with other cognitive domains (for a review, see Harvey, 2019). While the cognitive subcomponents of executive function are difficult to parse, further research should investigate the structural and functional associations of the cerebellum within each subcomponent of executive function.

## CONCLUSION

It is well-established that the prefrontal cortex and the cerebellum are highly age-sensitive brain regions. The current study was designed to isolate the cerebello-thalamo-cortical (CTC) tract fibers via diffusion tractography, assess the effects of age on its white matter integrity, and the role of the CTC in working memory and executive function performance. We report significant age-accelerated worsening of radial, axial, and mean diffusivities and that higher CTC diffusivities were associated with poorer executive function performance in an age-dependent manner, beginning around late middle-age. These findings underscore the role of the cerebellum and the CTC tract in higher order cognition, and begin to bridge the effects of two highly age vulnerable brain regions, the prefrontal cortex and the cerebellum.

## Appendix

**Table 1:** Regression model parameters for linear and quadratic age on a.) MD, b.) RD, and c.) AD, controlling for sex and years of education.

| Variable | B | *t* | *p* | *sr*<sup>2</sup> |
| --- | --- | --- | --- | --- |
| Sex | -0.011 | -1.911 | 0.057 | 0.01 |
| Education (years) | -0.001 | -0.639 | 0.524 | <0.01 |
| Age (centered) | -0.059 | -3.524 | <0.001 | 0.05 |
| Age<sup>2</sup> (centered) | 0.080 | 4.813 | <0.001 | 0.09 |

b.) Radial Diffusivity (RD)
| Variable | B | *t* | *p* | *sr*<sup>2</sup> |
| --- | --- | --- | --- | --- |
| Sex | -0.014 | -2.431 | 0.016 | 0.02 |
| Education (years) | -0.002 | -1.288 | 0.120 | <0.01 |
| Age (centered) | -0.032 | -1.878 | 0.062 | 0.01 |
| Age<sup>2</sup> (centered) | 0.052 | 2.980 | <0.003 | 0.04 |

c.) Axial Diffusivity (AD)
| Variable | B | *t* | *p* | *sr*<sup>2</sup> |
| --- | --- | --- | --- | --- |
| Sex | -0.004 | -0.605 | 0.546 | <0.01 |
| Education (years) | 0.001 | 0.653 | 0.514 | <0.01 |
| Age (centered) | -0.111 | -5.807 | <0.001 | 0.11 |
| Age<sup>2</sup> (centered) | 0.138 | 7.184 | <0.001 | 0.17 |

*Note.* *sr*<sup>2</sup> – Semi-partial correlations describe the unique contribution of each regression term on the dependent variable after controlling for all other terms in the model.; B = unstandardized beta weights

**Table 2:** Regression model parameters for diffusivity metric (a. MD, b. RD, and c. AD) on executive function composite score, controlling for sex and years of education.

| **Variable** | **B** | ***t*** | ***p*** | ***sr*<sup>2</sup>** |
| --- | --- | --- | --- | --- |
| Sex | -0.287 | -2.915 | 0.004 | 0.03 |
| Education (years) | 0.034 | 1.721 | 0.087 | 0.01 |
| MD | -1.654 | -1.273 | 0.204 | <0.01 |
| Age (centered) | 1.813 | 2.028 | 0.044 | 0.01 |
| MD*Age (centered) | -2.808 | -2.455 | 0.015 | 0.02 |

b.) RD and Executive Function
| **Variable** | **B** | ***t*** | ***p*** | ***sr*<sup>2</sup>** |
| --- | --- | --- | --- | --- |
| Sex | -0.296 | -2.971 | 0.003 | 0.03 |
| Education (years) | 0.032 | 1.652 | 0.100 | <0.01 |
| RD | -1.925 | -1.504 | 0.134 | <0.01 |
| Age (centered) | 1.038 | 1.561 | 0.120 | <0.01 |
| RD*Age(centered) | -2.498 | -2.145 | 0.033 | 0.02 |

c.) AD and Executive Function
| **Variable** | **B** | ***t*** | ***p*** | ***sr*<sup>2</sup>** |
| --- | --- | --- | --- | --- |
| Sex | -0.268 | -2.739 | 0.007 | 0.03 |
| Education (years) | 0.036 | 1.821 | 0.070 | 0.01 |
| AD | -0.802 | -0.756 | 0.451 | <0.01 |
| Age (centered) | 2.747 | 2.438 | 0.016 | 0.02 |
| AD*Age(centered) | -2.604 | -2.785 | 0.006 | 0.03 |

*Note.* *sr*<sup>2</sup> – Semi-partial correlations describe the unique contribution of each regression term on the dependent variable after controlling for all other terms in the model.; B = unstandardized beta weights

**Table 3:** Regression model parameters for diffusivity metric (a. MD, b. RD, and c. AD) on working memory composite score, controlling for sex and years of education.

| **Variable** | **B** | ***t*** | ***p*** | ***sr*<sup>2</sup>** |
| --- | --- | --- | --- | --- |
| Sex | -0.207 | -2.026 | 0.044 | 0.02 |
| Education (years) | 0.048 | 2.376 | 0.019 | 0.02 |
| MD | -0.960 | 1.349 | 0.478 | <0.01 |
| Age (centered) | -0.600 | -0.646 | 0.519 | <0.01 |
| MD*Age (centered) | 0.277 | 0.233 | 0.816 | <0.01 |

b.) RD and Working Memory
| **Variable** | **B** | ***t*** | ***p*** | ***sr*<sup>2</sup>** |
| --- | --- | --- | --- | --- |
| Sex | -0.216 | -2.100 | 0.038 | 0.02 |
| Education (years) | 0.047 | 2.311 | 0.022 | 0.02 |
| RD | -1.409 | -1.067 | 0.288 | <0.01 |
| Age (centered) | -0.606 | -0.883 | 0.379 | <0.01 |
| RD*Age(centered) | 0.402 | 0.334 | 0.739 | <0.01 |

c.) AD and Working Memory
| **Variable** | **B** | ***t*** | ***p*** | ***sr*<sup>2</sup>** |
| --- | --- | --- | --- | --- |
| Sex | -0.198 | -1.944 | 0.053 | 0.01 |
| Education (years) | 0.049 | 2.388 | 0.018 | 0.02 |
| AD | 0.031 | 0.028 | 0.978 | <0.01 |
| Age (centered) | -0.322 | -0.275 | 0.784 | <0.01 |
| AD*Age(centered) | -0.066 | -0.067 | 0.946 | <0.01 |

*Note.* *sr*<sup>2</sup> – Semi-partial correlations describe the unique contribution of each regression term on the dependent variable after controlling for all other terms in the model.; B = unstandardized beta weights

